# Prediction of rapid kidney function decline using machine learning combining blood biomarkers and electronic health record data

**DOI:** 10.1101/587774

**Authors:** Girish N. Nadkarni, Fergus Fleming, James R. McCullough, Kinsuk Chauhan, Divya A. Verghese, John C. He, John Quackenbush, Joseph V. Bonventre, Barbara Murphy, Chirag R. Parikh, Michael Donovan, Steven G. Coca

## Abstract

**Introduction:** Individuals with type 2 diabetes (T2DM) or the *APOL1* high-risk genotype (*APOL1*) are at increased risk of rapid kidney function decline (RKFD) as compared to the general population. Plasma biomarkers representing inflammatory and kidney injury pathways have been validated as predictive of kidney disease progression in several studies. In addition, routine clinical data in the electronic health record (EHR) may also be utilized for predictive purposes. The application of machine learning to integrate biomarkers with clinical data may lead to improved identification of RKFD.

**Methods:** We selected two subpopulations of high-risk individuals: T2DM (n=871) and *APOL1* high risk genotype of African Ancestry (n=498), with a baseline eGFR ≥ 45 ml/min/1.73 m^2^ from the Mount Sinai Bio*Me* Biobank. Plasma levels of tumor necrosis factor 1/2 (TNFR1/2), and kidney injury molecule-1 (KIM-1) were measured and a series of supervised machine learning approaches including random forest (RF) were employed to combine the biomarker data with longitudinal clinical variables. The primary objective was to accurately predict RKFD (eGFR decline of ≥ 5 ml/min/1.73 m^2^/year) based on an algorithm-produced score and probability cutoffs, with results compared to standard of care.

**Results:** In 871 participants with T2DM, the mean age was 61 years, baseline estimated glomerular filtration rate (eGFR) was 74 ml/min/1.73 m^2^, and median UACR was 13 mg/g. The median follow-up was 4.7 years from the baseline specimen collection with additional retrospective data available for a median of 2.3 years prior to plasma collection. In the 498 African Ancestry patients with high-risk *APOL1* genotype, the median age was 56 years, median baseline eGFR was 83 ml/min/1.73 m^2^,and median UACR was 11 mg/g. The median follow-up was 4.7 years and there was additional retrospective data available for 3.1 years prior to plasma collection. Overall, 19% with T2DM, and 9% of the APOL1 high-risk genotype experienced RKFD. After evaluation of three supervised algorithms: random forest (RF), support vector machine (SVM), and Cox survival, the RF model was selected. In the training and test sets respectively, the RF model had an AUC of 0.82 (95% CI, 0.81-0.83) and 0.80 (95% CI, 0.78-0.82) in T2DM, and an AUC of 0.85 (95% CI, 0.84-0.87) and 0.80 (95% CI, 0.73-0.86) for the *APOL1* high-risk group. The combined RF model outperformed standard clinical variables in both patient populations. Discrimination was comparable in two sensitivity analyses: 1) Using only data from ≤ 1 year prior to baseline biomarker measurement and 2) In individuals with eGFR ≤60 and/or albuminuria at baseline. The distribution of RFKD probability varied in the two populations. In patients with T2DM, the RKFD score stratified 18%, 49%, and 33% of patients to high-, intermediate-, and low-probability strata, respectively, with a PPV of 53% in the high-probability group and an NPV of 97% in the low-probability group. By comparison, in the APOL1 high-risk genotype, the RKFD score stratified 7%, 23%, and 70% of patients to high-, intermediate-, and low-probability strata, respectively, with a 46% PPV in the high-probability and an NPV of 98% NPV in the low-probability group.

**Conclusions:** In patients with T2DM or of African Ancestry with the high-risk *APOL1* genotype, a RF model derived from plasma biomarkers and longitudinal EHR data significantly improved prediction of rapid kidney function decline over standard clinical models. With further validation, this approach may be valuable in aiding clinicians in identifying patients who would benefit most from early and more aggressive follow-up to mitigate kidney disease progression.

## INTRODUCTION

Type 2 diabetes (T2DM) is being recognized as an epidemic worldwide. One of the most common complications of T2DM is diabetic kidney disease (DKD) which develops in over one third of T2DM cases. DKD is the single largest cause of end-stage renal disease (ESRD), accounts for 44% of incident ESRD patients and is a major independent risk factor for other complications including coronary artery disease, stroke, and retinopathy. In addition, the rates of ESRD are higher among persons with African ancestry (AA) compared to European Americans (EAs) across all baseline estimated glomerular filtration rate (eGFR) levels.^1,2^ Genetic admixture studies demonstrated that two distinct alleles in the Apolipoprotein L1 (*APOL1*) gene on chromosome 22 confer substantially increased risk for a number of kidney diseases in AA, including focal segmental glomerulosclerosis, human immunodeficiency virus-associated nephropathy, and hypertension-attributable kidney disease. The *APOL1* high-risk genotypes (i.e., two copies of the *APOL1* renal risk variants; G1/G1; G2/G2 or G1/G2) are associated with increased ESRD risk, chronic kidney disease (CKD) progression,^3^ eGFR decline,^4^ and incident CKD.^5^ Thus, ancestry differences in *APOL1* risk prevalence could partly explain disparities in kidney disease between AAs and EAs.

Even though these populations are on average higher risk than the general population, prediction of who will have rapid kidney function decline (RKFD) is challenging.^6,7^ Currently, the prevalent standard for ESRD risk prediction in CKD Stages 3-5 is the kidney failure risk equation (KFRE), where clinical variables (including age, sex, estimated glomerular filtration rate [eGFR] and urine albumin creatinine ratio [UACR]) are assigned standard weights for a recursive score calculation. However, the KFRE has not been validated in individuals with normal kidney function at baseline, where arguably intervention would have the most impact.^8^

Several blood and urine biomarkers have been investigated to aide with the prediction of incident and progressive CKD.^9^ Three of the most extensively studied biomarkers and most associated with CKD progression are soluble tumor necrosis factor 1/2 (TNFR1/2), and plasma kidney injury molecule-1 (KIM-1).^10–18^ While these markers have uniformly shown independent associations with CKD and CKD progression along with specific clinical variables such as eGFR and UACR, implementation of accurate models which combine clinical data with these plasma biomarkers to predict CKD progression is lacking.

Widespread use of electronic health records (EHR) provides the potential to leverage thousands of longitudinal clinical features for prediction of events. Standard statistical approaches are inadequate to fully leverage the EHR due to thousands of features, unaligned nature of data, and correlation structure.^3^ However, contemporary supervised machine learning approaches have the analytical model building capacity to combine both biomarkers and longitudinal EHR data for improved prediction.

In the current study, we utilized retrospectively collected plasma samples linked to longitudinal clinical data from the Icahn School of Medicine at Mount Sinai (ISMMS) Bio*Me* Biobank to examine the ability of supervised machine learning algorithms to predict rapid kidney function decline.

## MATERIALS AND METHODS

### The Bio*Me* Biobank at Icahn School of Medicine at Mount Sinai (ISMMS)

The Bio*Me* Biobank at ISMMS is an Institutional Review Board (IRB)-approved plasma and DNA biorepository protocol which includes consented access to the patients’ electronic medical record (EMR) from a diverse local community in New York City.^10,11^ Bio*Me* operations were initiated in 2007 and are fully integrated in clinical care processes, including direct recruitment from over 30 broadly selected clinical sites by dedicated recruiters. For the purpose of this study, we selected two subpopulations: 1) Bio*Me* participants with T2DM, an eGFR between 45 and 90 ml/min/1.73 m2 at the time of Bio*Me* enrollment, and at least ≥3 years of follow up data in the EHR; 2) Bio*Me* participants with *APOL1* high risk genotype, eGFR > 30 ml/min/1.73m^2^ at the time of Bio*Me* enrollment and at least ≥ 3 years of follow up data in the EHR.

### Ascertainment and Definition of the kidney endpoint

We determined eGFR using the CKD-EPI creatinine equation, calculated median values per 3-month period of follow up and utilized these for outcome ascertainment. We defined the primary outcome as “rapid kidney function decline (RKFD)” as an eGFR decline of ≥ 5 ml/min/1.73 m^2^/year with a minimum of 3 values after the baseline date.^19–24^

### Ascertainment of clinical variables in Bio*Me* Biobank

Sex and AA race were obtained from an enrollment questionnaire administered to Bio*Me* participants. Clinical data were extracted for all continuous variables (eGFR, hemoglobin A1c, urine protein or albumin to creatinine ratios) at baseline from the EHR with concurrent time stamps. We defined the baseline period as 1 year before the Bio*Me* enrollment date. Body mass indices (BMI) were calculated as the ratio between weight and the square of height in kg/m^2^. Hypertension and T2DM status at baseline were determined using the Electronic Medical Records and Genomics (eMERGE) Network phenotyping algorithms.^16^ Cardiovascular disease and heart failure were determined by a validated algorithm from the Electronic Medical Records and Genomics (eMERGE) network and ICD-9/10 codes respectively. We considered a participant to be on an angiotensin converting enzyme-inhibitor (ACE-i) or angiotensin receptor blocker (ARB) if they had a concurrent prescription during the Bio*Me* enrollment. We considered baseline values as the median of all values in the 1 year period immediately prior to the enrollment date We calculated follow up time from Bio*Me* enrollment date to latest visit in the EHR.

### Biospecimens Storage and Analytes Measurement

Plasma specimens were collected on the day of enrollment into Bio*Me*. The plasma samples were stored at − 80°C. Biomarkers were measured using the Mesoscale platform (Meso Scale Diagnostics, Gaithersburg, Maryland, USA), which employs proprietary electrochemiluminescence detection methods combined with patterned arrays to allow for multiplexing of assays. The intra- and inter-assay coefficient of variation (CV) for the quality control samples were 3.5%, 3.9%, and 4.5%, and 12.4%, 10.8%, and 7.7%, for TNFR1, TNFR2, and KIM1, respectively. The average lower limit of detection (LOD) obtained from multiple runs was 12.5 pg/ml for TNFR1, 7.8 pg/ml for TNFR2, and 9.0 pg/ml for KIM1 (**Supplementary Tables 1 and 2**). The laboratory personnel performing the biomarker assays were blinded to clinical information about the participants.

### Statistical Analysis

We expressed descriptive results for the participants’ baseline characteristics and biomarkers via means and standard deviations, or for skewed variables, medians and interquartile ranges. Statistical comparisons between groups were performed by paired t-tests for data that were normally distributed, Wilcoxon tests for skewed continuous data, and McNemar’s test for categorical data.

We evaluated three supervised learning algorithms on the combined dataset of biomarker values and longitudinal clinical variables (truncated at the date of biomarker measurement). For the model, we considered two data inputs i. Biomarker measurements ii.Structured EHR features including laboratory values, diagnosis/procedure codes, socio-demographics, medications and healthcare encounter history. We then created meta-features from these existing structured variables including maximum, minimum, median, variability and change over time to account for the longitudinal aspect and repeated nature.

The three supervised algorithms evaluated were random forest (RF), conditional inference forest (CIF) and supervised vector machine (SVM). For all algorithms, the data were randomly split to create a training group representing 80%, and a testing group representing 20% of the population. Additionally, we conducted 10-fold cross-validation on all models. Algorithm method selection was based on AUC performance using DeLong’s test.

We then conducted further iterations of the model by tuning the individual hyperparameters. For example, Hyperparameter 1, is the number of decision trees in the forest, hyperparameter 2, the number of variables randomly selected for splitting at each node and hyperparameter 3, minimum size of terminal nodes. The final model was chosen which had the best AUC compared to previous iterations. As a comparison to current standards, we compared the comprehensive algorithm derived using biomarkers and clinical variables (combined RF model) to three base models via likelihood ratio tests: 1) the Tangri 4-Variable kidney failure risk equation (KFRE), 2) biomarker only model (logistic regression) and 3) only EHR features using RF model.

We conducted two sensitivity analyses: 1. Using only data from ≤ 1 year prior to biomarker measurement (i.e., “contemporary data”) and 2. In individuals with existing CKD (eGFR ≤ 60 ml/min/1.73 m^2^ and/or albuminuria at baseline). We compared all differences between AUCs using a DeLong’s test for comparisons.

Finally, we calculated a RKFD probability score from 0-100 for the machine learning algorithm based on the predicted individual probabilities of having rapid kidney function decline and scaled it using log transformation. We defined low, intermediate and high probability strata using population cutoffs and calculated sensitivity, specificity and positive/negative predicted values (PPV/NPV). Goodness of fit statistics were used to assess calibration of the RFKD score vs. the observed outcomes. All analyses were performed with R software (www.rproject.org)

## RESULTS

### Baseline Characteristics of Cohorts

#### Patients with T2DM (n=871)

The median age was 60 years, 507 (58%) were female, and median eGFR was 68 ml/min/1.73 m^2^ (**Table 1**). The most common comorbidities were hypertension (93%), coronary heart disease (50%), and heart failure (22%). The majority (77%) were on ACE inhibitors or angiotensin receptor blockers (ARBs). Patient characteristics including events between the training and test cohorts were balanced (**Supplementary Table 3**).

**Table 1.** Clinical and Characteristics

|  | Type 2 DM (n=871) | APOL1 (n=498) |
| --- | --- | --- |
| **Clinical Characteristics** |  |  |
| Age in years, Median (IQR) | 60 [53 - 66] | 56 [46-66] |
| Female, n (%) | 507 (58.2) | 337 (67.6) |
| Race |  |  |
| White | 62 (7.1) | 0 |
| African-American | 346 (39.7) | 498 (100) |
| Hispanic/Latino | 412 (47.3) | 0 |
| Other | 51 (5.9) | 0 |
| Body Mass Index in kg/m<sup>2</sup>, Median [IQR] | 30.9 [26.6 - 36.1] | 30.5 [26.2-35.8] |
| Hypertension, n (%) | 813 (93.3) | 220 (44.2) |
| Coronary Artery Disease, n (%) | 432 (49.6) | 39 (7.8) |
| Heart Failure, n (%) | 192 (22.0) | 15 (3) |
| Systolic BP in mm Hg, Median [IQR] | 131.9 [123.7 - 143.4] | 129 [117-140.5] |
| Diastolic BP in mm Hg, Median [IQR] | 73.4 [68.7 - 79.3] | 77 [69.5-84.5] |
| Mean Arterial Pressure in mm Hg, Median [IQR] | 92.8 [88.1 - 99.9] | 93.7 [86-102] |
| Follow up Time in years, Median [IQR] | 4.5 [3.3 - 6.1] | 5.9 [3.9-7.1] |
| **Laboratory Characteristics** |  |  |
| Baseline eGFR, Median [IQR] | 68.4 [55.3 - 80.0] | 83.3 [68.9-99.4] |
| Baseline UACR, Median [IQR] | 13.0 [4.0 - 66.3] | 11 [4.5-55] |
| Baseline Hemoglobin A1C, Median [IQR] | 7.0 [6.2 - 8.7] | 5.9 [5.5-6.4] |
| **Medications** |  |  |
| ACE/ARB, n (%) | 675 (77.5) | 122 (24.5) |
| **Plasma Biomarker Concentrations** |  |  |
| TNFR1, in pg/ml, Median [IQR] | 6057.0 [4764.9 - 8224.4] | 2465 [1988-3266] |
| TNFR2, in pg/ml, Median [IQR] | 6914.2 [5332.9 - 9832.9] | 4215 [3234-5654] |
| KIM1, in pg/ml, Median [IQR] | 323.3 [196.8 - 592.1] | 154 [96-269] |

#### Patients with APOL1 High-Risk Genotype (n=498)

Median age was 56 years, 337 (67.6%) were female, and median eGFR was 83.3 ml/min/1.73 m^2^ (**Table 1**). The prevalence of comorbidities were much lower than compared to the cohort with T2DM, i.e. hypertension (44%), coronary heart disease (8%), and heart failure (3%). Patient characteristics including events between the training and test cohorts were comparable (**Supplementary Table 4**).

### Rapid Kidney Function Decline (RKFD) Endpoints

For participants with T2DM, 164 of the 871 (18.8%) experienced rapid kidney function decline over a median follow-up of 4.6 (IQR 3.4-5.6) years. In participants with *APOL1* high-risk genotype, 45 of the 498 (9%) experienced rapid kidney function decline over a median follow up of 5.9 (IQR 3.9-7.1) years.

### Machine learning models for prediction of rapid kidney function decline

In patients with T2DM, the RF model had an AUC=0.82, compared to an AUC=0.81 for SVM, and an AUC of 0.78 for Cox survival for predicting RKFD. In AA patients, the AUC for the RF model was 0.85, compared to AUC of 0.78 for SVM, and 0.83 for Cox survival [p<0.05 for comparisons]. Based upon these results and a knowledge base that RF performs better with data structures like EHR, RF models were chosen for further analysis, and all further results are presented using RF.

For the patients with T2DM, the training AUC for the combined RF model to predict RKFD was 0.82 (95% CI 0.80-0.90) on 10-fold cross validation and was 0.80 (95% CI 0.75-0.80) in test. By comparison, the KFRE for RKFD yielded an AUC of 0.57 (95% CI 0.56-0.58), the biomarker only model had an AUC of 0.76 (95% CI 0.72-0.79), and an optimized clinical model using RF (incorporating EHR data without biomarkers) produced an AUC of 0.74 (95% CI 0.73-0.76); **Figure 1A**). The top ten data features contributing to performance of the combined RF model were the three plasma biomarkers (TNFR1, TNFR2 and KIM1) and laboratory values or vital signs (either baseline or change over time) that are linked to kidney disease. The *P* value of the Hosmer-Lemeshow goodness-of-fit test for the combined RF model was 0.20, indicating there was no significant difference between the predicted and observed outcomes (**Supplementary Figure 1**).

**Figure 1A.**
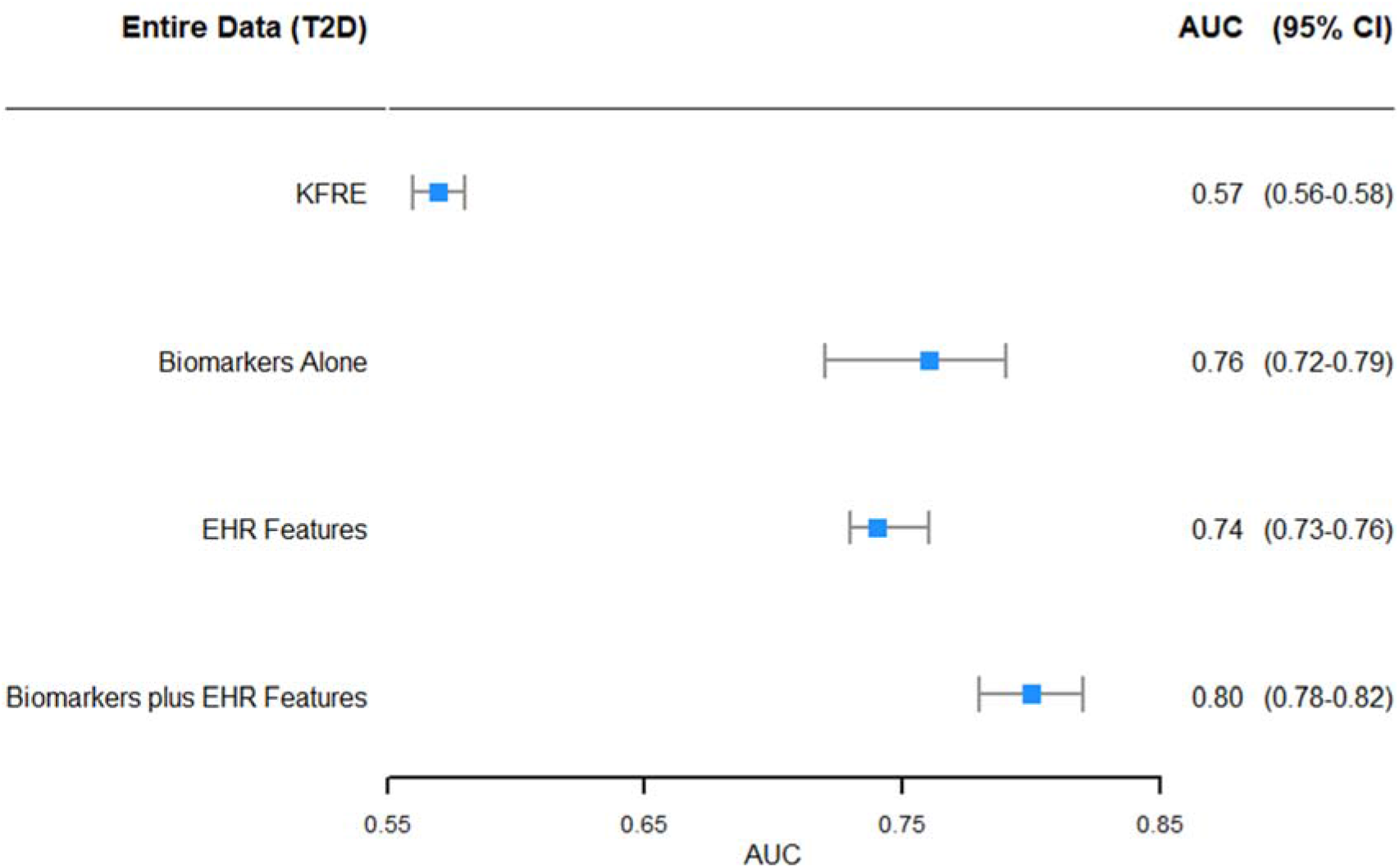
Comparison of AUCs Using Increasing Data Inputs in type 2 DM

For the patients with *APOL1* high risk genotype, the AUC of the RF was 0.85 (95% CI 0.84-0.87) on 10-fold cross validation in the training set and was 0.80 (95% CI 0.73-0.86) in the testing set. Discrimination of KFRE for RKFD had an AUC of 0.48 (95% CI 0.45-0.50), the biomarker model had an AUC of 0.71 (95% CI 0.64-0.78), and the optimized clinical model using RF (incorporating EHR data without biomarkers) had an AUC of 0.77 (95% CI 0.70-0.85; **Figure 1B**). The top ten data features contributing to the model performance in the AAs with *APOL1* high-risk genotype were the three plasma biomarkers (TNFR1, TNFR2 and KIM1) and and laboratory values or vital signs (either baseline or change over time) that are linked to kidney disease. The *P* value of the Hosmer-Lemeshow goodness-of-fit test was 0.41, indicating there was no significant difference between the predicted and observed outcomes (**Supplementary Figure 1B**).

**Figure 1B.**
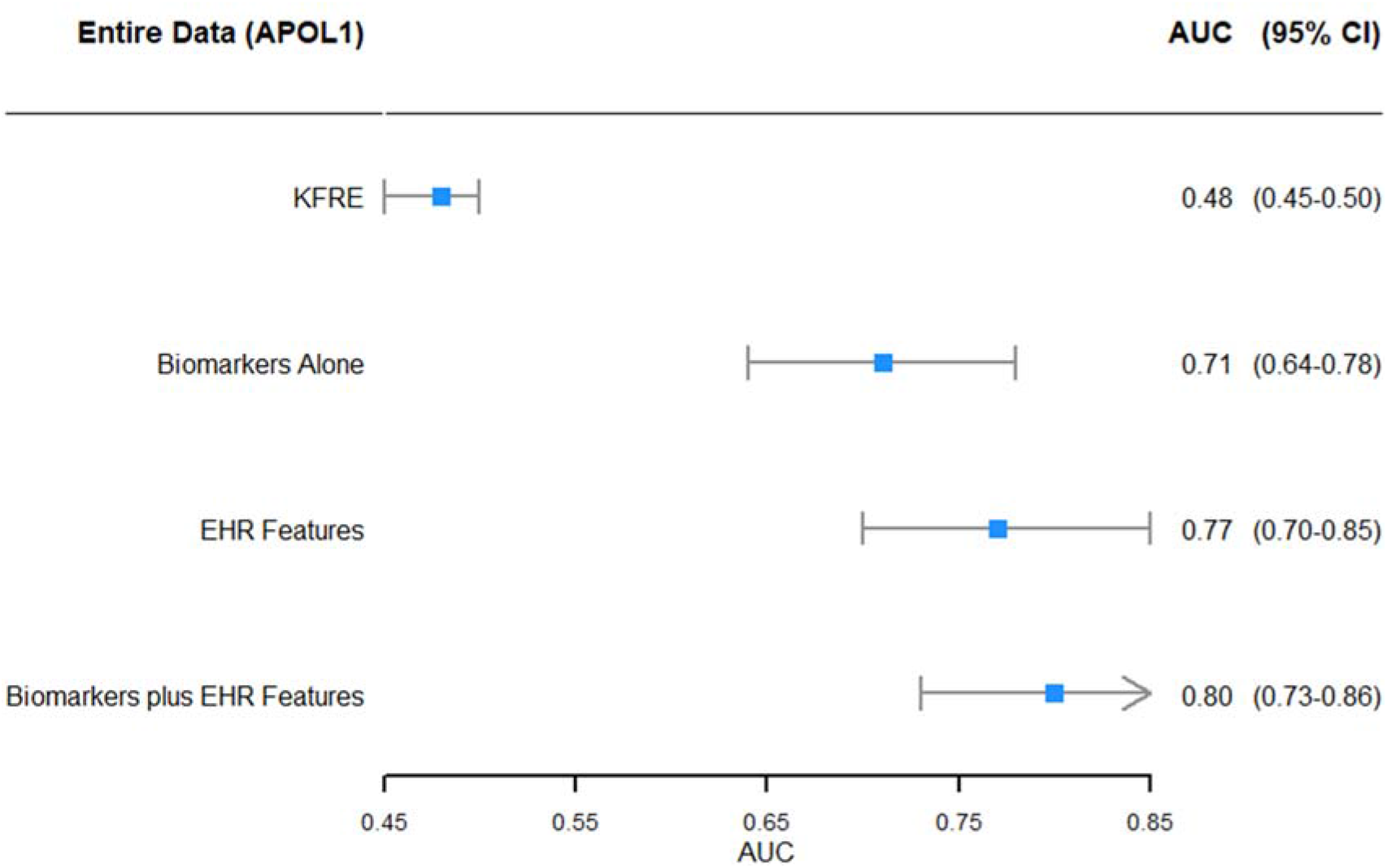
Comparison of AUCs Using Increasing Data Inputs in APOL1

### Sensitivity Analyses: Contemporary Data and Prevalent CKD

Using contemporary data only (data within 1 year prior to enrollment and biomarker measurement), the discriminatory performance of the combined RF model in both the T2DM (**Figure 2A**) and *APOL1* high-risk genotype were similar to the overall cohort results (**Figure 2B**).

**Figure 2A.**
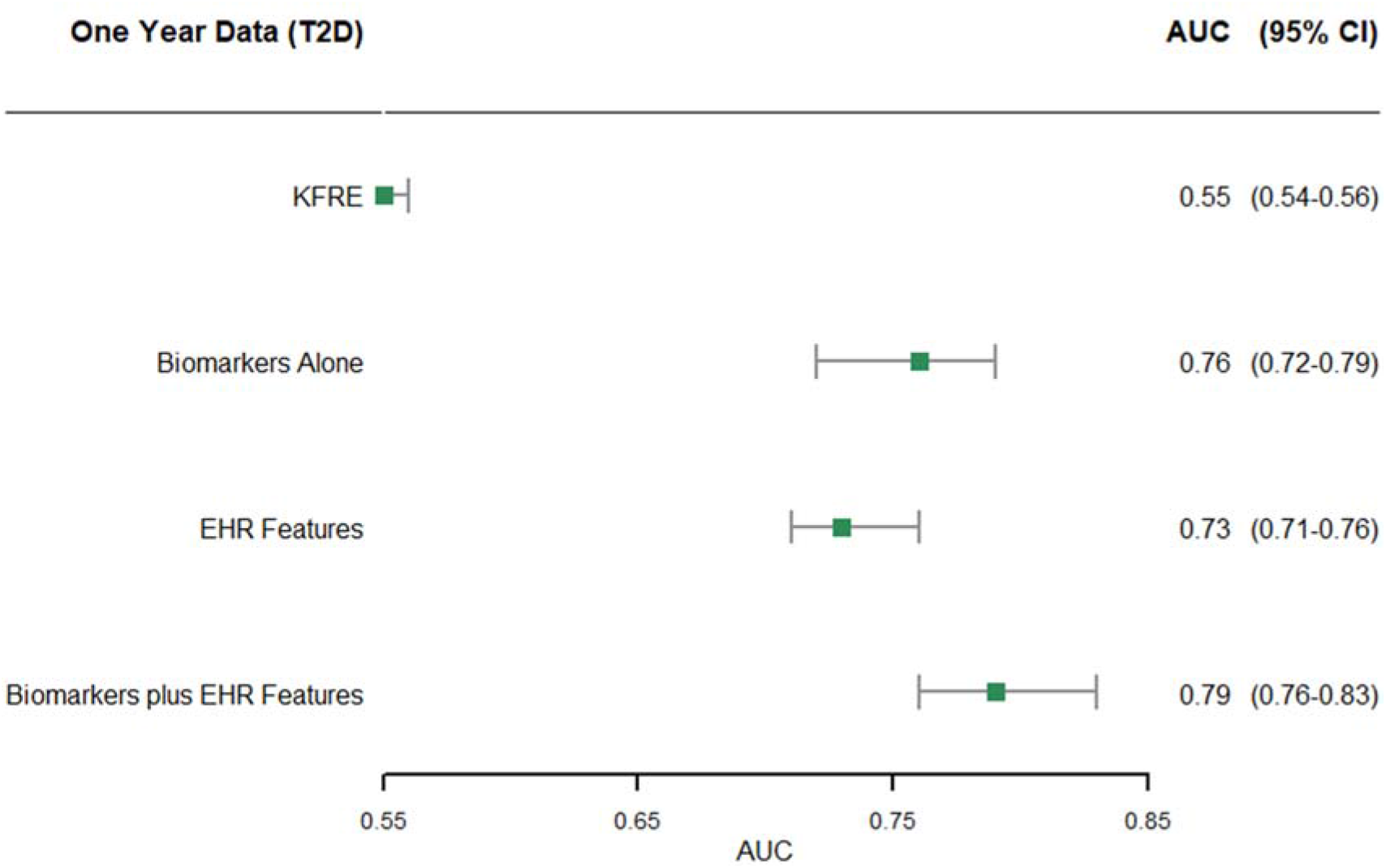
Comparison of AUCs Using Increasing Data Inputs using data from ≤1 year prior to sample collection (Contemporary Data) in Type 2 DM

**Figure 2B.**
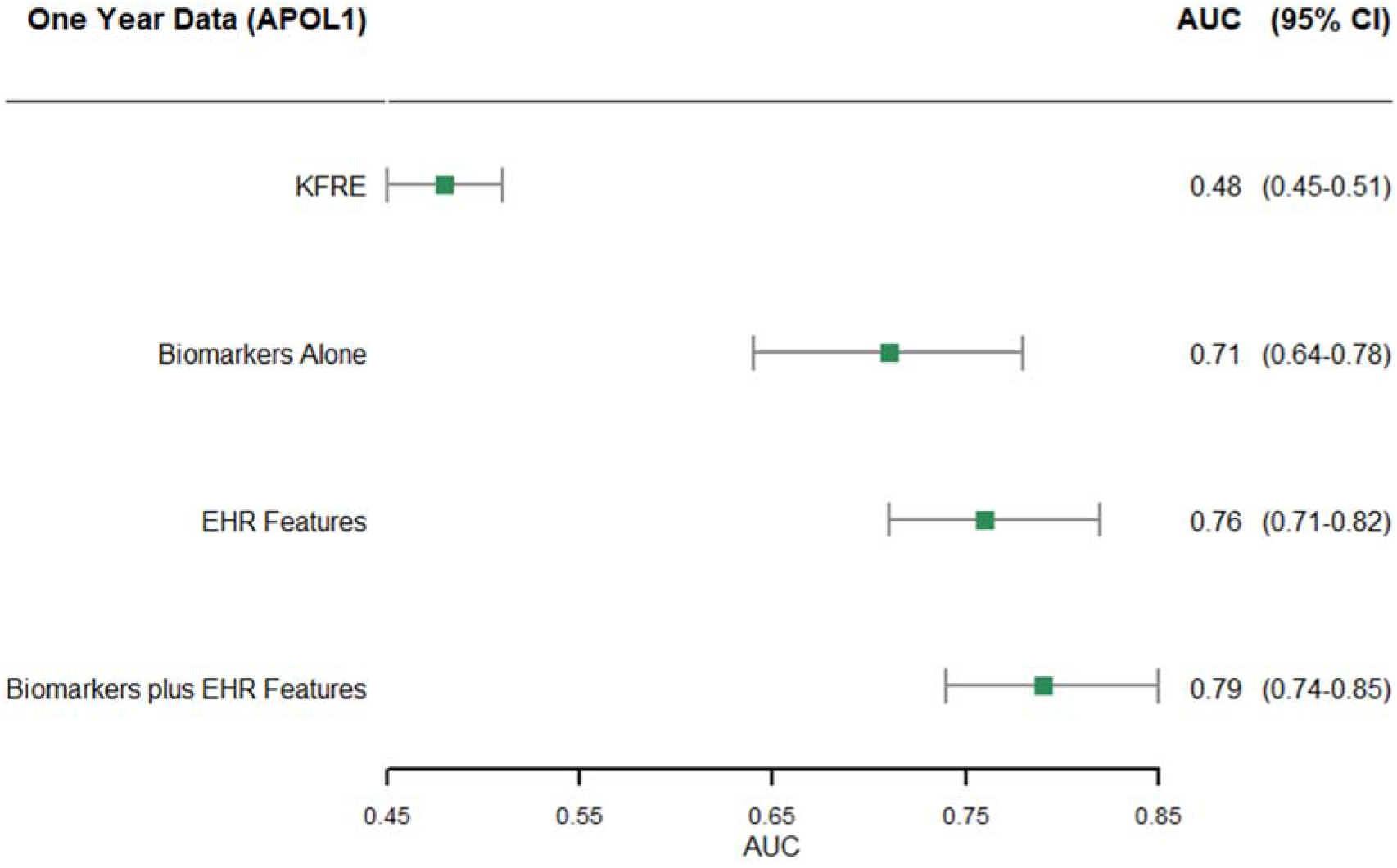
Comparison of AUCs Using Increasing Data Inputs using data from ≤1 year prior to sample collection (Contemporary Data) in APOL1

In a subset of theT2DM cohort with CKD, (eGFR ≤ 60 ml/min/1.73 m^2^ and/or albuminuria at baseline) (n=366), 21.5% experienced RKFD. Results for the combined RF model and comparison to the KFRE, biomarkers alone, and the clinical models were similar to the overall cohort results (**Figure 3A**).

**Figure 3A.**
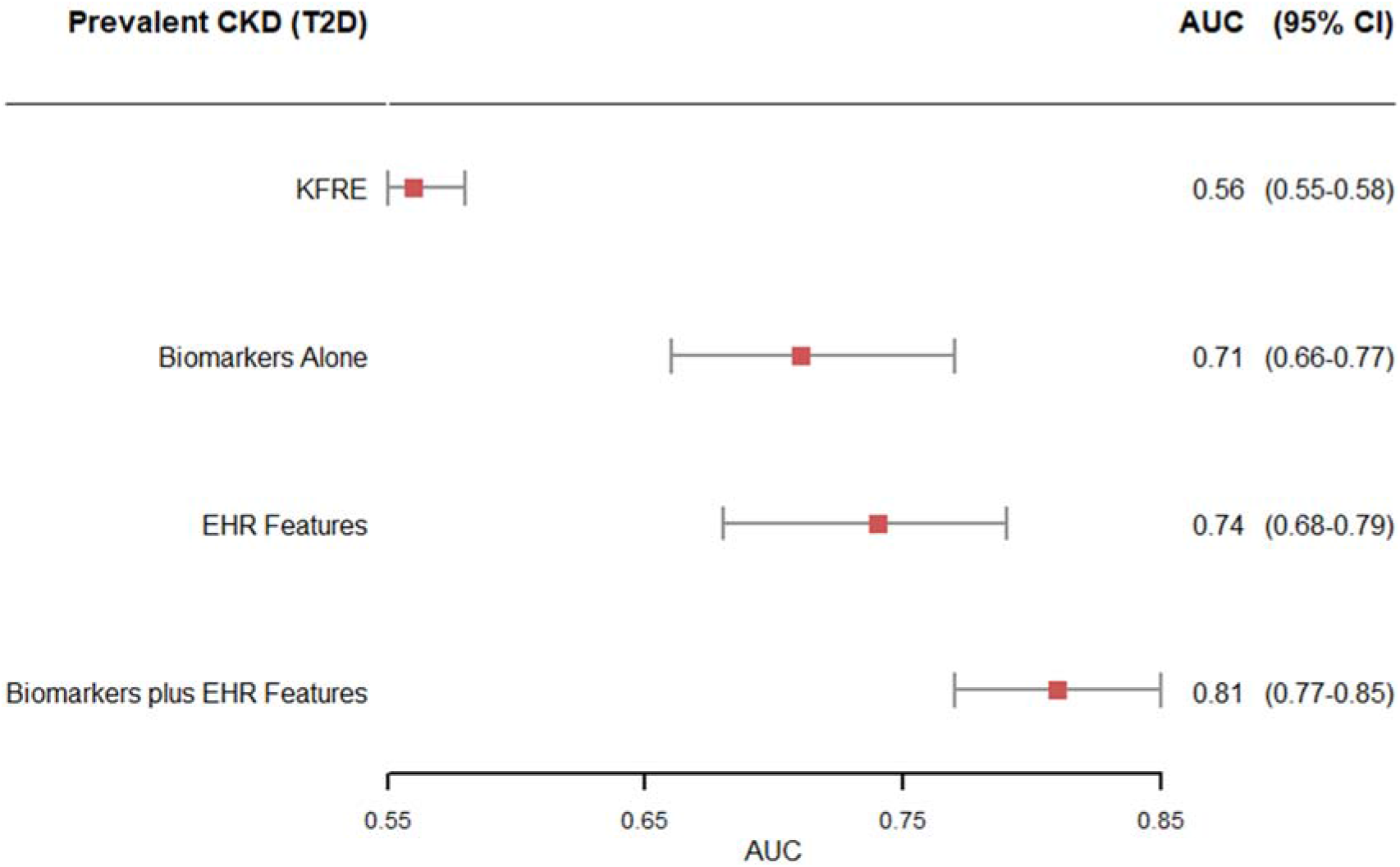
Comparison of AUCs Using Increasing Data Inputs in patients with prevalent CKD in Type 2 DM

In patients with *APOL1* high risk genotype, 112 patients had baseline prevalent CKD, of which 9% experienced RKFD. In this subgroup, the RF model had an AUC of 0.88, which outperformed the KFRE (AUC 0.49), and the biomarker only model (AUC 0.73), but was similar to the performance for an optimized RF model using only clinical features without biomarkers (AUC 0.87; **Figure 3B**).

**Figure 3B.**
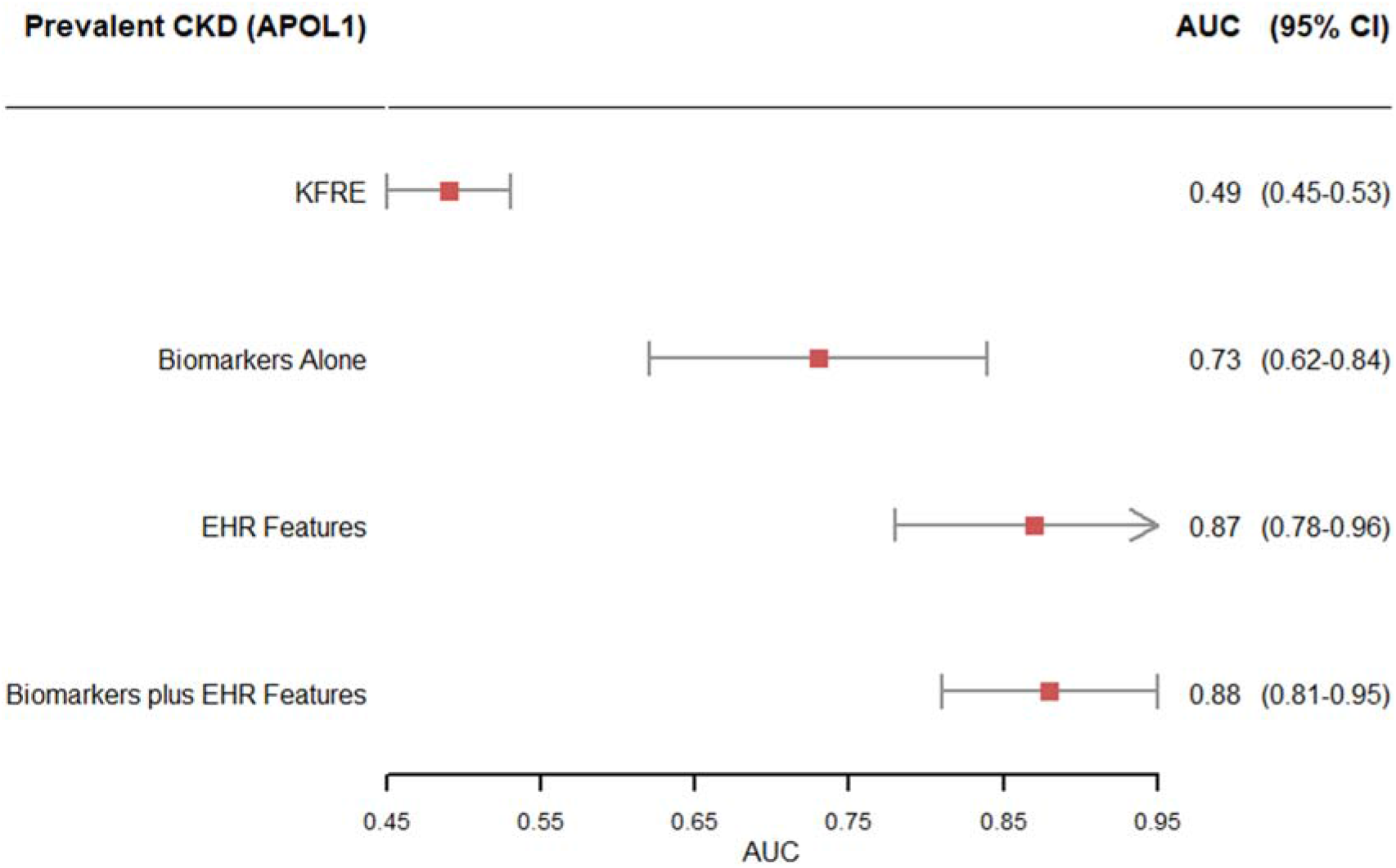
Comparison of AUCs Using Increasing Data Inputs in patients with prevalent CKD in APOL1

### RFKD Score for rapid kidney function decline (RKFD)

The combined RF models with biomarkers and clinical variables were used to generate cutoffs for low, intermediate and high predicted probability for RFKD (high, intermediate and low) and used to calculate the sensitivity, specificity, PPV and NPV for each pre-defined cutoff. In patients with T2DM, the RFKD Score stratified 18%, 49%, and 33% of patients to high-, intermediate-, and low-probability score strata, with a PPV in the high-probability group of 53% and a 97% NPV in the low-probability group (**Table 2, Figure 4A**). For the *APOL1* high-risk genotype cohort, the RFKD Score stratified 7%, 23%, and 70% of patients to high-, intermediate-, and low-probability groups, with a PPV in the high-probability group of 46% in the high group and a 98% NPV in the low-probability group. (**Table 2, Figure 4B**).

**Table 2.**
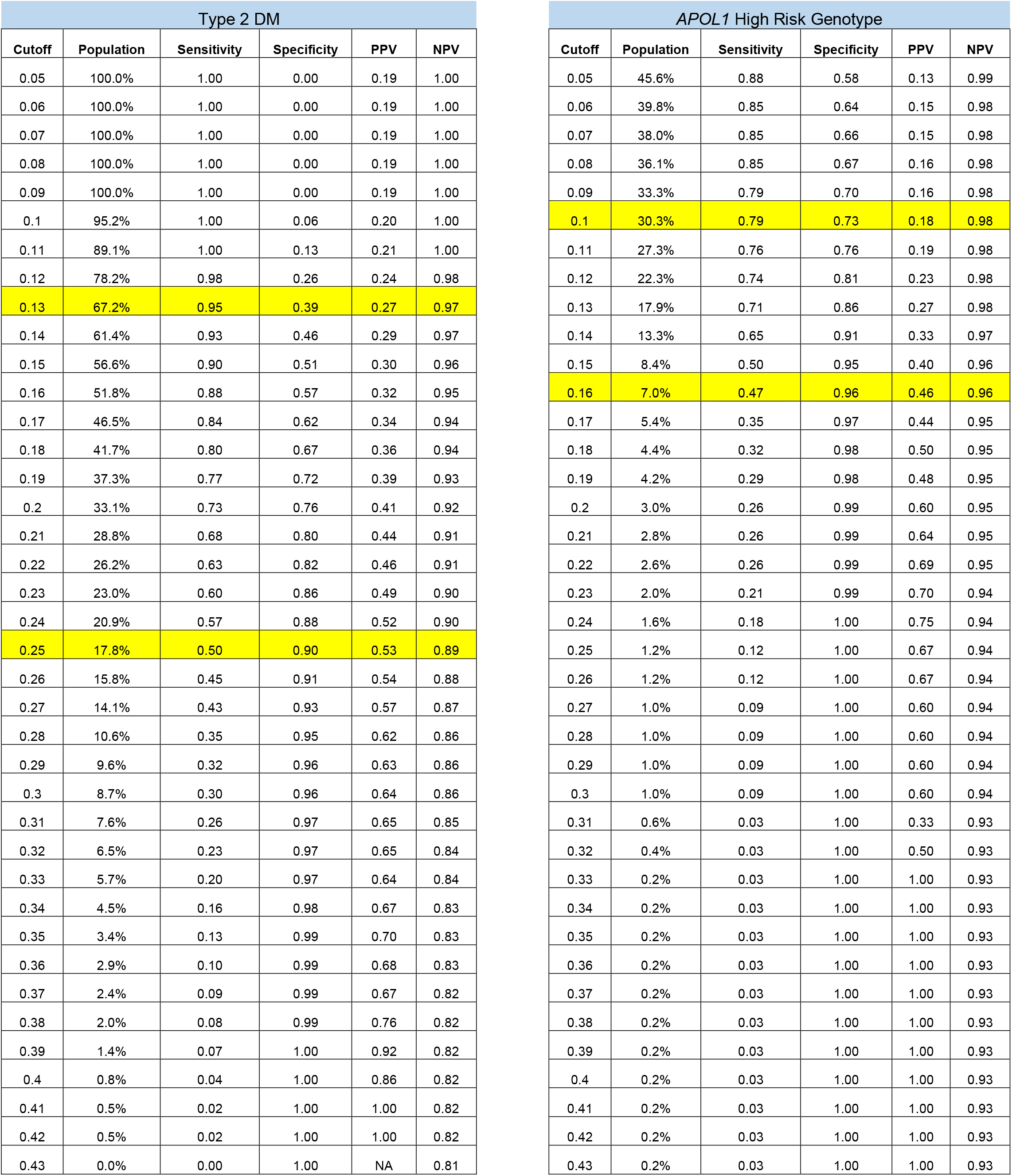
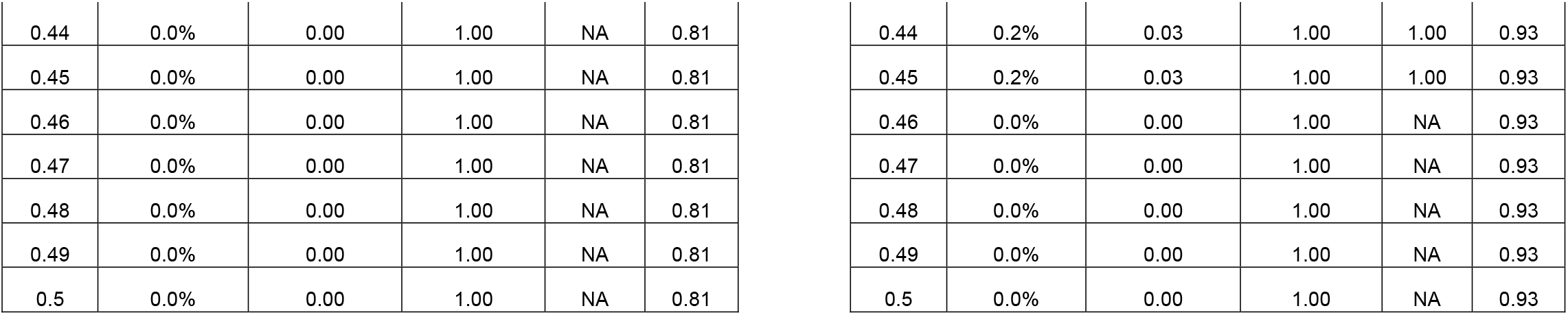
Combined ML Prediction Model Thresholds for RKFD with Sensitivity, Specificity, PPV and NPV for Type 2 DM and *APOL1* High-Risk Populations

**Figure 4A.**
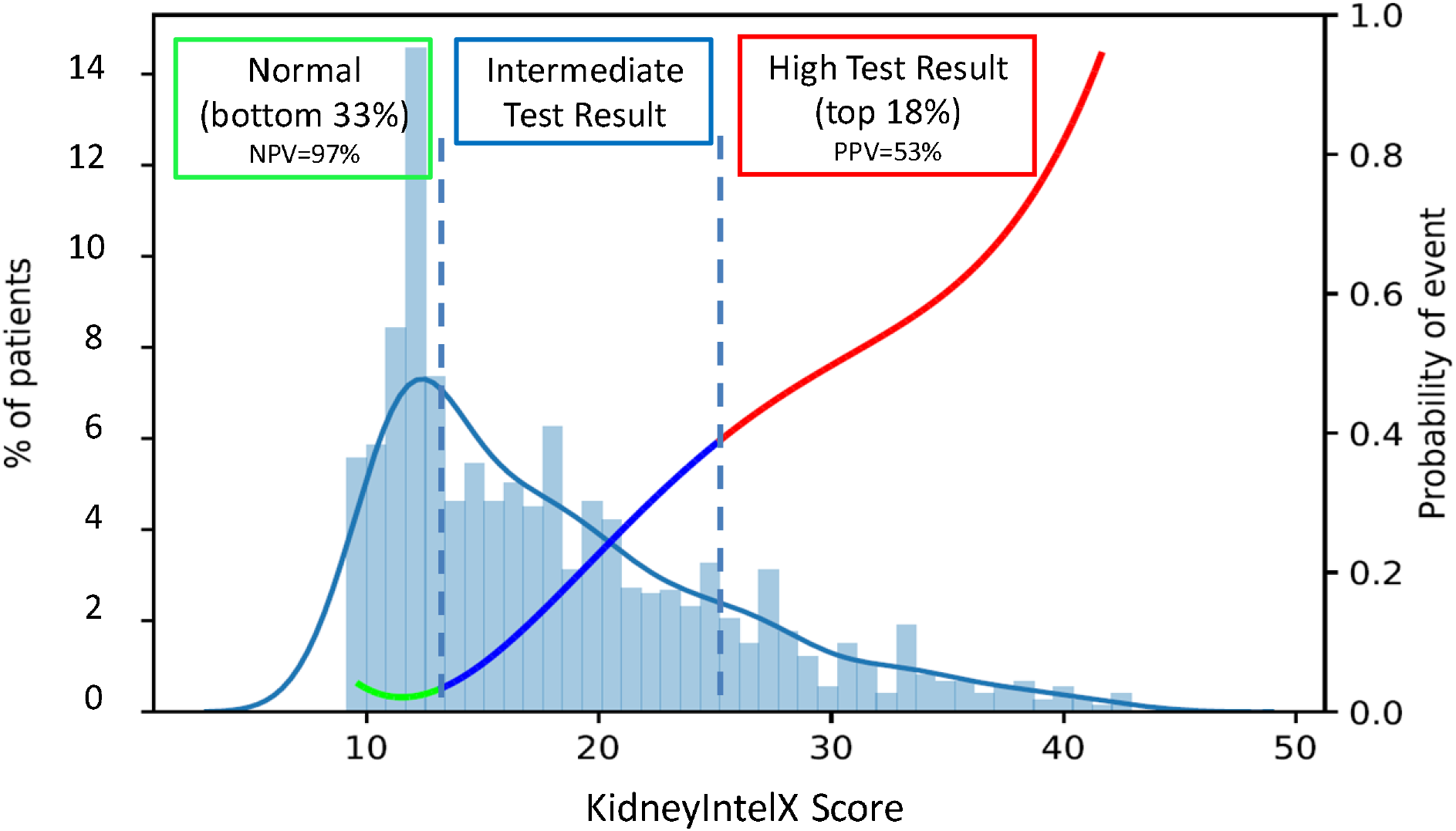
Distribution of RKFD Scores and Probability of Rapid Kidney Function Decline by Continuous and Categorical Strata in Type 2 DM. The blue histogram bars represent the proportion of patients with type 2 DM in each strata of the RKFD score. The green, blue, and red line represents the smoothed probability of experiencing rapid kidney function decline.

**Figure 4B.**
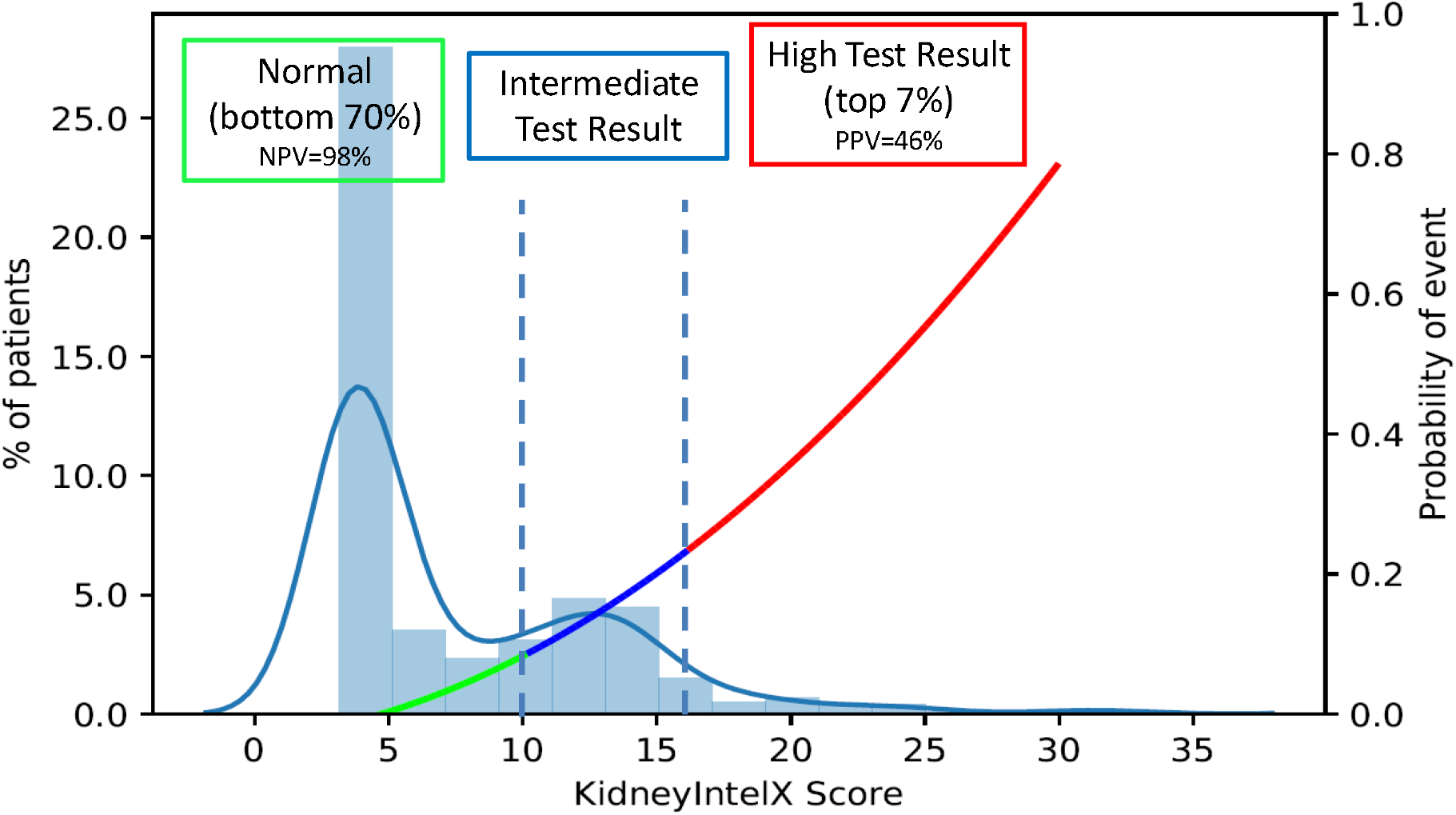
Distribution of RKFD Scores and Probability of Rapid Kidney Function Decline by Continuous and Categorical Strata in Patients with APOL1 High-Risk Genotype. The blue histogram bars represent the proportion of patients with *APOL1* High-risk Genotype in each strata of the RKFD score. The green, blue, and red line represents the smoothed probability of experiencing rapid kidney function decline.

## DISCUSSION

Utilizing two large cohorts of patients (T2DM and AAs with the high-risk *APOL1* genotype), both linked to the EHR and with retrospectively banked plasma samples, we developed a random forest supervised machine learning algorithm combining extant, longitudinal EHR data and three previously validated plasma biomarkers to predict rapid kidney function decline (RKFD). The combined RF model outperformed standard clinical metrics and the current standards for prediction of RKFD, including albuminuria and other clinical variables.^8,25^ The ability to identify a distinct group of patients with the RKFD score with a PPV of approximately 50% for RKFD should allow for more appropriate patient management including referral to a nephrologist, improved awareness of overall kidney health, and guidance towards more targeted, intensive therapies to slow RKFD. The demonstrated PPV in the high RFKD score strata represented a 3-5 fold improvement over current standard of care and the observed baseline event rate in the two populations.

CKD is a complex, common problem challenging modern healthcare. In the absence of specific therapies to cure CKD, early identification of patients more likely to experience RKFD are paramount. Early identification would help in allocation of limited resources as well as implementation or intensification of proven interventions to slow kidney function decline. In real world practice, the prediction of RKFD in patients with T2DM and *APOL1* high-risk genotype are challenging, particularly in patients with largely preserved kidney function. There are two major problems contributing to the difficulties in early identification and prediction: 1) serum creatinine/eGFR and urine albumin creatinine are relatively insensitive and non-specific biomarkers for kidney function decline, with significant fluctuations and variability in early stages of CKD, and 2) the prevalent standard includes recursive scores incorporating only a single (baseline) value of a selected predictive feature and does not include the comprehensive, longitudinal data that is present in the EHR.

Recently, several biomarkers representing the pathways of injury and inflammation have been the subject of an intense research focus in CKD. Among these, three biomarkers have sufficient evidence and appear appropriate for clinical efficacy and implementation. These biomarkers are soluble tumor necrosis factor receptor 1 and 2 (sTNFR1/2) and plasma kidney injury molecule-1 (KIM-1) and each have been extensively validated in multiple studies including patients with^10–13,15–17,26^ and without T2DM^26,27^ as well as patients with the high risk *APOL1* genotype.^28^ The individual biomarkers have added significant improvement to classical clinical metrics for RKFD prediction, and the combination of all three, perhaps because of different pathophysiologic pathways, appears to be synergistic.^28^ We have demonstrated that combining these biomarkers with clinical information using advanced machine learning techniques can significantly improve discrimination and prediction of RKFD.

Along with biomarkers that can be measured during routine clinical encounters, as demonstrated in the current BioMe collection, the longitudinal EHR data present in most large health care systems can be combined with these biomarker values for optimal clinical prediction. We have shown previously that the addition of temporal, longitudinal data using supervised machine learning significantly outperforms ‘baseline’ clinical models and also has significant utility for subtyping disease trajectories.^29,30^ Thus, we hypothesized that combining biomarker information and extant longitudinal EHR data would significantly improve the prediction of the subsequent RKFD, as demonstrated in this confirmational study.

This integrated approach has several, near-term clinical implications, especially when linked to clinical decision support (CDS) and embedded care pathways within the EMR. For example, patients with a high RFKD score, who will have a predicted probability of > 50% of experiencing rapid kidney function decline should be referred to a nephrologist, which has been shown to be associated with improvement in outcomes.^31^ In addition, referral to a dietician and delivery of educational materials regarding the importance and consequences of CKD can be provided to the high RFKD score patients to help increase awareness and facilitate motivation for changes in lifestyles and behavior. Finally, the optimization of medical therapy including renin-angiotensin aldosterone system inhibitors, statins for cardiovascular risk management, and intensification of antihypertensive medication to meet guideline recommended blood pressure targets can be pursued. The application of sodium glucose transporter (SGLT)-2 inhibitors might also be advantageous in the high RFKD score group with T2DM given the recent data on robust renoprotection with these agents.^32–34^ On the other hand, patients with a low predicted probability could be clinically managed by their primary care provider and have standard of care treatment with scheduled monitoring of their RFKD scores. Finally, patients with an intermediate RFKD score would be recommended for standard of care and retesting longitudinally. Such patients may demonstrate score shifts based on behavior, clinical parameters and treatments change over time with appropriate clinical actions as necessary. This overall approach would not only have benefits for individual patient outcomes but also at a health system and population level, where there is uncertainty about which patients to refer to a limited number of subspecialists.

Our study does have some limitations. Although we utilized a large, multiethnic cohort in this initial confirmation study, extended validation in geographically diverse cohorts is needed. Secondly, data structures and relationships change over time due to adjusted practice patterns and thus the algorithm may not perform similarly if the full complement of longitudinal data are not available. We conducted a sensitivity analysis with only one year of data available prior to the baseline, and the loss of performance was minimal, however, this should be evaluated further in additional validation cohorts. Finally, this study does not address implementation or utility. Therefore, in addition to further validation, a randomized controlled trial to assess whether providing this score to providers and patients through EMR systems and perhaps linking to CDS/clinical pathways improves the process of care or patient outcomes.

In conclusion, we have demonstrated that using machine learning techniques (random forest models) to combine longitudinal EHR information with three novel, validated plasma biomarkers, improved prediction of RKFD over standard clinical models. With the advent of advanced high performance computing, validated biomarkers and integrated EHRs, the paradigm of implementation of the RKFD score should be tested for integration into routine patient care for improved outcomes.

## Author Contributions

Drs. Nadkarni and Coca had full access to all of the data in the study and take responsibility for the integrity of the data and the accuracy of the data analysis.

*Concept and design:* Nadkarni, Donovan, Coca

*Acquisition, analysis, or interpretation of data:* Nadkarni, Fleming, Donovan, Coca

*Drafting of the manuscript:* Nadkarni, Coca

*Critical revision of the manuscript for important intellectual content:* All authors

*Statistical analysis:* Nadkarni, Fleming, Coca

*Administrative, technical, or material support:* Chauhan, Verghese, McCullough

*Supervision:* Nadkarni, Fleming, Murphy, Donovan, Coca

---

**Supplementary Figure 1A.**
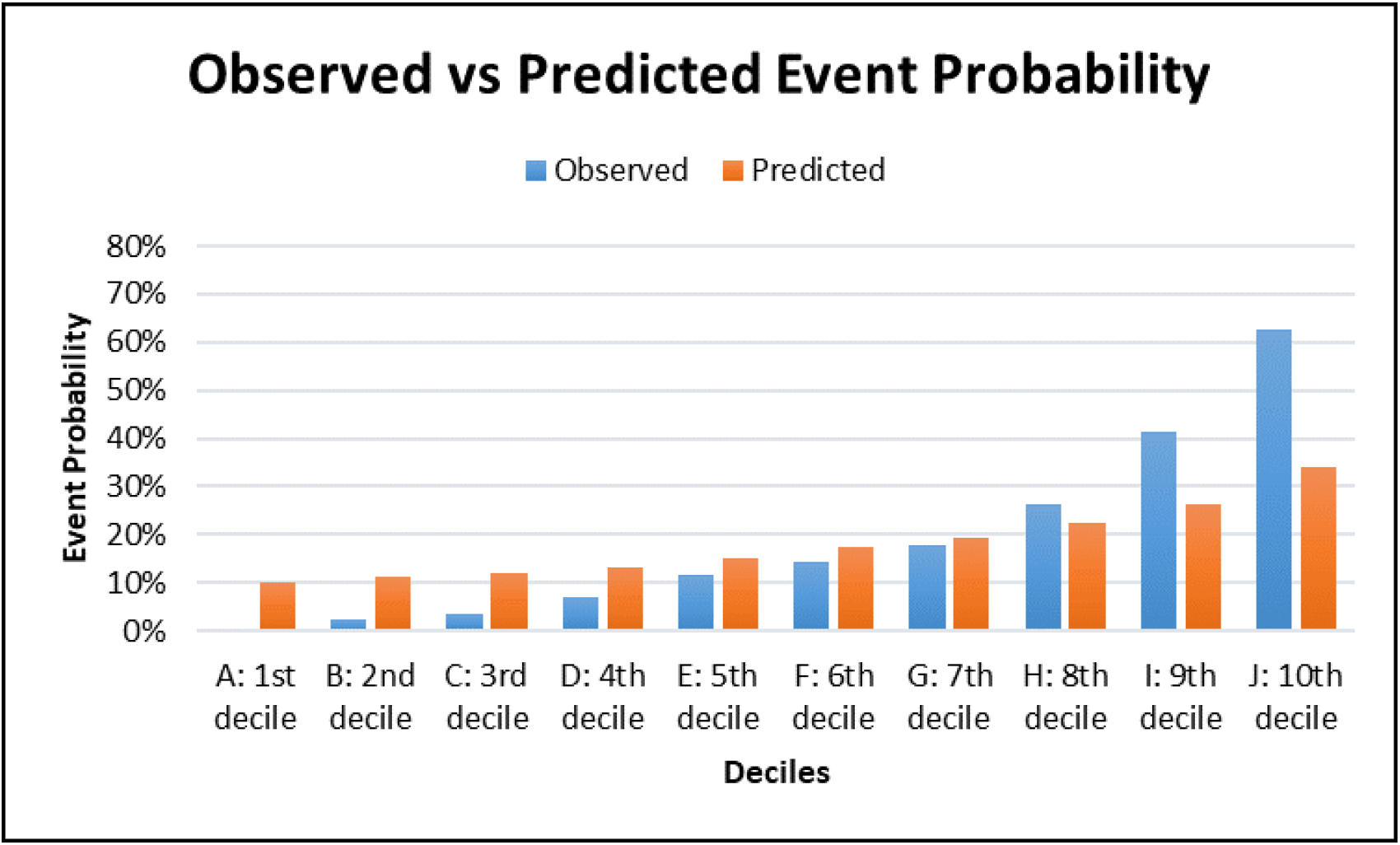
Observed vs. Expected (calibration plot) in Patients with Type 2 DM

**Supplementary Figure 1B.**
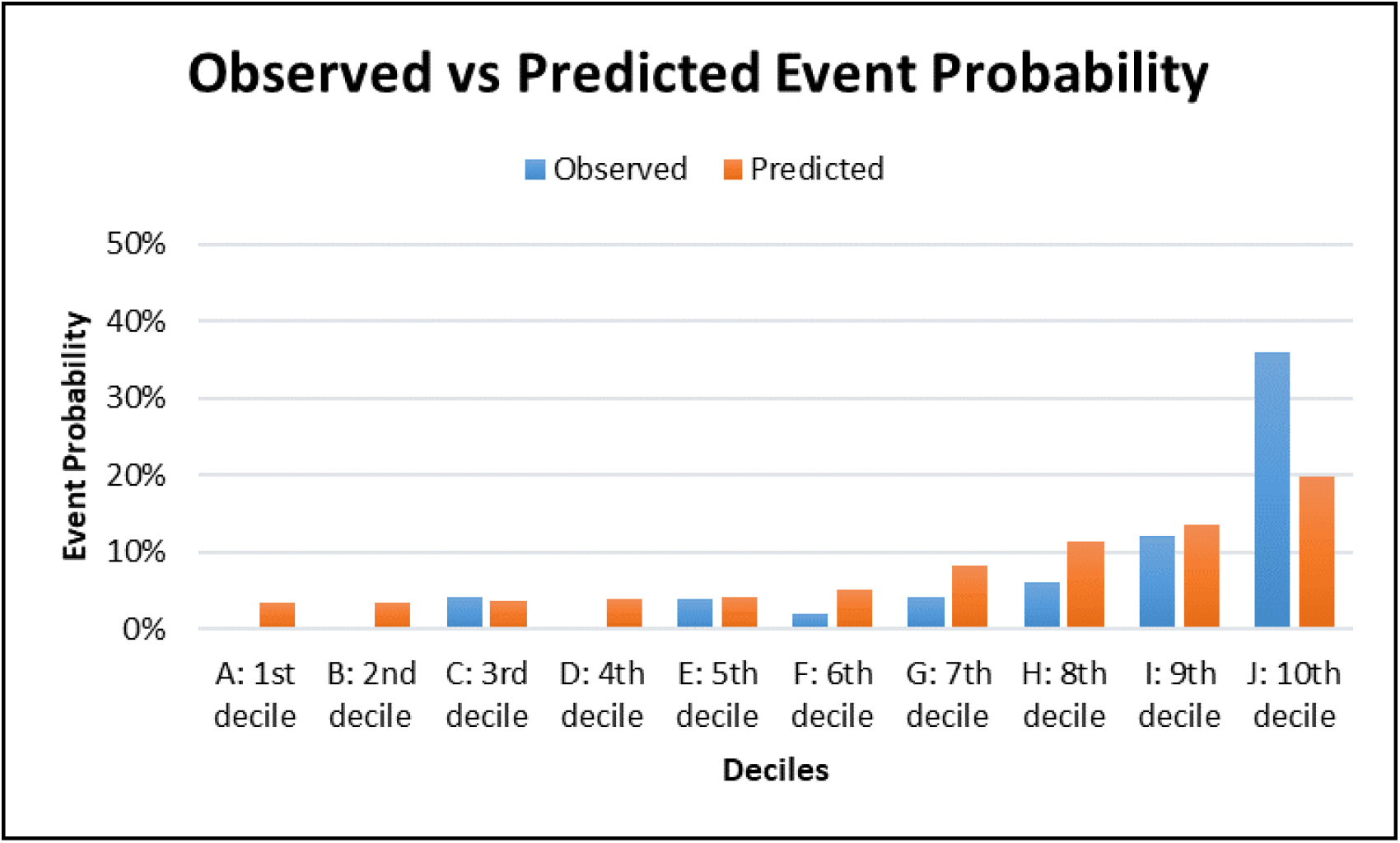
Observed vs. Expected (calibration plot) in Patients with *APOL1* High-Risk Genotype

**Supplementary Table 1.** Intra- and Inter-assay CV% and Lower Limit of Detection

| Marker | Intra-assay CV % (low) | Intra- assay CV% (high) | Inter-assay CV% (low) | Inter-assay CV% (high) | LLOD (pg/ml) |
| --- | --- | --- | --- | --- | --- |
| Kim-1 | 4.5 | 2.9 | 14.9 | 8.7 | 2.1 |
| TNF-R1 | 3.4 | 4.7 | 15.0 | 14.3 | 7.9 |
| TNF-R2 | 4.4 | 2.8 | 9.9 | 10.1 | 0.3 |

**Supplementary Table 2.** Spike and Recovery and Linearity of Dilution

|  | KIM-1 |  |  | TNF-R1 |  |  | TNF-R2 |  |  |
| --- | --- | --- | --- | --- | --- | --- | --- | --- | --- |
|  | Conc (pg/ml) | CV% | % Recovery | Conc (pg/ml) | CV% | % Recovery | Conc (pg/ml) | CV% | % Recovery |
| High Spike | 1773.0 | 2.6 | 86.7 | 3868.7 | 0.04 | 102.9 | 3765.8 | 1.7 | 97.1 |
| Medium Spike | 448.6 | 1.4 | 82.4 | 3039.0 | 0.59 | 101.0 | 2987.1 | 0.08 | 95.4 |
| Low Spike | 165.6 | 7.4 | 97.8 | 2799.0 | 2.32 | 99.2 | 2944.3 | 5.3 | 100.1 |
| No Spike | 44.4 | 3.5 | 100.0 | 2758.2 | 5.00 | 100.0 | 2880.4 | 2.6 | 100.0 |

**Supplementary Table 3.** Key Characteristics in Training and Test Datasets in T2DM Cohort

|  | Train (n= 696) | Test (n= 175) |
| --- | --- | --- |
| Age in years, Median [IQR] | 60 [53 - 67] | 59 [51 - 66] |
| Female, n (%) | 411 (59%) | 96 (54.9 %) |
| Race, n (%) |  |  |
| African American | 272 (39%) | 74 (42.2%) |
| European American | 48 (6.9%) | 14 (8 %) |
| Hispanic Latino | 337 (48.4%) | 75 (42.9 %) |
| Others | 39 (5.6) | 1 (0.57 %) |
| Biomarkers, Median [IQR] |  |  |
| TNFR1 | 6025 [4765 - 8105] | 6250 [4812 - 9097] |
| TNFR2 | 6825 [5262 - 9763] | 7429 [5712 - 10312] |
| KIM1 | 313.11 [196.96 - 562.36] | 379.3 [201.6 - 722.5] |
| eGFR in ml/min, Median [IQR] | 70.04 [55.35 - 82.12] | 68.27 [55.8 - 82] |
| Systolic BP in mm Hg, Median [IQR] | 132 [120 - 146] | 128 [120 - 140] |
| Diastolic BP in mm Hg, Median [IQR] | 75 [67 - 83] | 74 [66 - 80] |
| RFKD events, n (%) | 125 (18%) | 31 (18%) |

**Supplementary Table 4.** Key Characteristics in Training and Test Datasets in *APOL1* High Risk Cohort

|  | **Train**<br>**(n= 398)** | **Test**<br>**(n= 100)** |
| --- | --- | --- |
| Age in years, Median [IQR] | 49 [39 - 59] | 52 [43 - 62] |
| Female, n (%) | 269 (67.6 %) | 68 (68 %) |
| Race, n (%) |  |  |
| African American | 377 (95) | 94 (94) |
| Hispanic Latino | 9 (2.3) | 4 (4) |
| Others | 12 (3) | 2 (2) |
| Biomarkers, Median [IQR] |  |  |
| TNFR1 | 2467.7 [1989.7 - 3203] | 2441 [1971 - 3397] |
| TNFR2 | 4254.8 [3231.9 - 5627.8] | 4180 [3256 - 5675] |
| KIM1 | 159.6 [96 - 275.5] | 141.3 [96.2 - 236.7] |
| eGFR in ml/min, Median [IQR] | 82 [68.3 - 98.5] | 82.4 [66.1 - 99.3] |
| Systolic BP in mm Hg, Median [IQR] | 130.2 [117 - 140] | 131 [118.0 - 143] |
| Diastolic BP in mm Hg, Median [IQR] | 76.8 [70 - 84.8] | 79.5 [70 - 86] |
| RFKD events, n (%) | 32 (8) | 7 (7) |

